# High-throughput microgel biofabrication via air-assisted co-axial jetting for cell encapsulation, 3D bioprinting, and scaffolding applications

**DOI:** 10.1101/2022.10.08.511435

**Authors:** Vaibhav Pal, Yogendra Pratap Singh, Deepak Gupta, Mecit Altan Alioglu, Momoka Nagamine, Myoung Hwan Kim, Ibrahim T. Ozbolat

**Author notes:** Corresponding author: Ibrahim T. Ozbolat. Authors contributed equally (Vaibhav Pal, Yogendra Pratap Singh).

## Abstract

Microgels have recently received widespread attention for their applications in a wide array of domains such as tissue engineering, regenerative medicine, and cell and tissue transplantation because of their properties like injectability, modularity, porosity, and the ability to be customized in terms of size, form, and mechanical properties. However, it is still challenging to mass produce microgels with diverse sizes and tunable properties. Herein, we developed an air-assisted co-axial device (ACAD) for continuous production of microgels in a high-throughput manner. To test its robustness, microgels of multiple hydrogels and their combination, including alginate (Alg), gelatin methacrylate (GelMA) and Alg-GelMA, were formed at a maximum production rate of 65,000 microgels per sec while retaining circularity and a size range of 50-500 μm based on varying air pressure levels. The ACAD platform allowed single and multiple cell encapsulation with around 75% efficiency. These microgels illustrated appealing rheological properties such as yield stress, viscosity, and shear modulus for bioprinting applications. Specifically, Alg microgels have the potential to be used as a sacrificial support bath while GelMA microgels have potential for direct extrusion both on their own or when loaded in a bulk GelMA hydrogel. Generated microgels showed high cell viability (>90%) and proliferation over 7 days with their increased interactions with cells, particularly for GelMA microgels. The developed strategy provides a facile and rapid approach without any complex or expensive consumables and accessories for scalable high-throughput microgel production for cell therapy, tissue regeneration and 3D bioprinting applications.

## 1. Introduction

Recent advances in tissue engineering have shown microgels to be a progressively used versatile class of materials, as scaffolds or drug/bioactive molecule delivery platforms [1]. They are crosslinked polymeric networks that offer unique properties of injectability, highly adaptable modular microstructure, along with biocompatibility and drug and cells encapsulation capabilities. Their extensive tunability has made them applicable in a wide range of tissue engineering applications [2,3]. Further, their performance for specific applications can be tuned by altering their physicochemical properties. Particularly, the size of microgels has a significant impact on their kinetic and thermodynamic properties as well as on their interactions with the surroundings [4]. Microgel size also directly influences the amount of drug or cells that can be encapsulated inside and later released. Therefore, high-throughput biofabrication of microgels with uniform size and shape is urgently needed to meet criteria for various tissue engineering, biofabrication and therapeutic applications.

With increased interest in microgels, various microgel fabrication techniques have been developed and advanced including micromolding, emulsion-based, shearing, and extrusion techniques [5–7]. The use of these techniques depends on the final application and compatibility of the biofabrication process with the hydrogel’s gelation mechanism. Recently, the microfluidic emulsion technique is particularly prevalent because of its capacity for microscale fluid manipulation [8,9]. This technique involves mixing two immiscible fluids—oil and a hydrogel precursor—to create droplets of the hydrogel precursor in oil using a flow-focusing device, typically a chip [10]. The ability to control the size and shape of the microgels, makes microfluidics the method of choice for many applications. However, these systems suffer from high manufacturing costs coupled with an escalating level of system design complexity, limited microgel production rate, and the lack of universality for including various solvents [11]. Therefore, a method for producing monodisperse microgels in a high-throughput, cost-effective and rapid manner is still desired. There have been numerous attempts to create microscale particles made of polyelectrolyte complexes. Herrero et al. developed a microencapsulation technology to produce sodium alginate particles (1–50 μm) using compressed air [12]. Wang et al. developed a miniature gas-liquid coaxial flow device using glass capillaries, aiming to produce sub-100-μm calcium-alginate microspheres [13]. These are emerging as versatile tools for aiding in the regeneration of damaged tissues.

Recently, much research has been focused on using microgels as a bioink for 3D bioprinting applications by improving the structural integrity and maintaining biological functions. This requires smooth extrusion of microgels as well as adequate mechanical strength to withhold 3D bioprinted constructs whilst preserving viability of encapsulated cells [2,3,14]. In one of the initial reports on microgel fabrication, Du et al. developed a bottom-up approach to control the assembly of cell-laden microgels to produce tissue constructs with adjustable complexity and microarchitecture. They used the thermodynamic tendency of multiphase liquid–liquid systems to minimize their contact surfaces to form poly(ethylene glycol) microgels [15]. Recently, alginate microgels have been used as a yield-stress support bath for aspiration-assisted bioprinting, which enabled the assembly of human mesenchymal stem cell spheroids for bone and cartilage tissue fabrication [16,17]. Concurrently, using a flow-focusing microfluidic device, core–shell gelatin methacrylate (GelMA) microgels have been produced [18]. The results showed biocompatibility of microgels by encapsulating liver cells in the cores, with the ability to coculture liver cells and vascular endothelial cells, indicating that core–shell architectures are suitable for heterocellular cultures. Utilizing the injectable shear thinning capabilities of microgels can help create highly customizable systems that can be used for cell and tissue (i.e., islet) delivery [19,20]. 3D Bioprinted microgels are emerging as promising platforms for generating tissue substitutes. To control the printability of the components and modulate desired physicochemical characteristics of the constructs more efficiently, development of bioinks based on jammed particles has been demonstrated [21]. Recently, Ouyang et al. developed a microgel-templated porogel bioink platform for controlled microporosity [22]. The addition of gelatin microgels significantly enhanced the printability of bioinks at various cell densities. The approach enables the engineering of highly tunable pores in cell-laden bioinks with controlled porosity and pore size; however, the fabrication process is conventional and low-throughput, and the removal of gelatin microgels by diffusion is a slow process.

Herein, we present a facile biofabrication method to produce microgels using air-assisted co-axial device (ACAD). The device setup had a concentric arrangement of two separate nozzles through which the polymer solution and air flowed (**Fig. 1**). To demonstrate the versatility of the process, microgels of multiple materials and their combination were fabricated including alginate (Alg), GelMA (GelMA), and alginate-GelMA (Alg-GelMA) microgels. The formed microgels were tested for multiple applications including their utilization for cell encapsulation, bioprinting (as a bioink for direct extrusion or as a support bath for embedded bioprinting), and scaffolding.

**Fig. 1:**
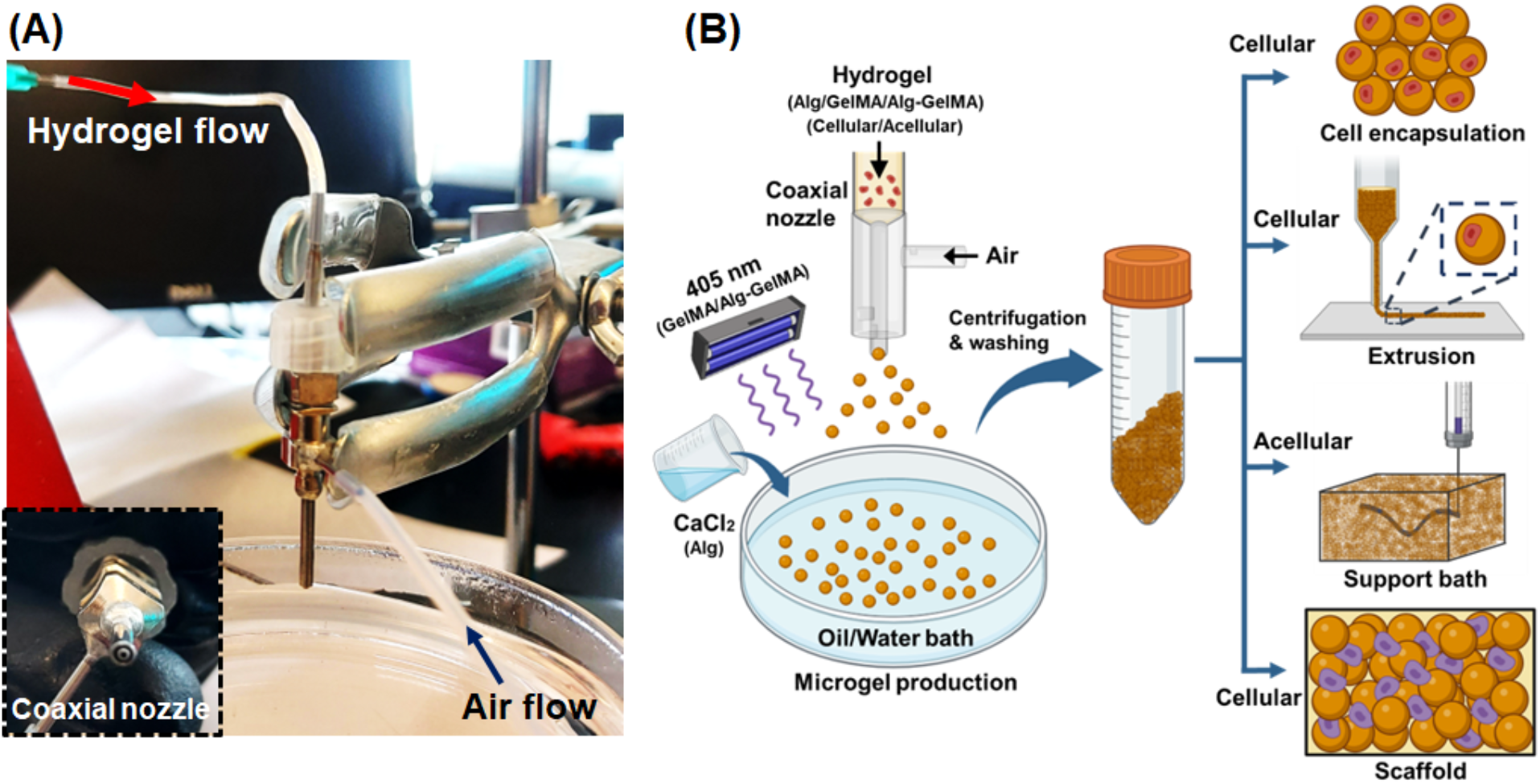
High-throughput fabrication of microgels. (A) The experimental setup, which involves a coaxial nozzle that was attached to an air pump for air flow and a syringe pump for hydrogel flow, where a collector bath was maintained below the coaxial nozzle. (B) The schematic diagram of microgel production from Alg, GelMA and Alg-GelMA hydrogels using the ACAD device. Microgels were collected, washed, and centrifuged for multiple applications including cell encapsulation, bioprinting via direct extrusion of cell encapsulated microgels or support bath for embedded bioprinting, and scaffolding.

## 2. Materials and Methods

### 2.1 Device Fabrication and Setup

The developed ACAD platform had four main components as shown in **Fig. 1A**. The first component was the coaxial nozzle system, where one nozzle was used for material flow and the other one for air flow. The second component was the air controller system (Ultimus I, Nordson EFD, East Providence, RI), which was used to regulate air flow. The air controller system was connected to the outer nozzle (15 G) of the coaxial assembly. The third component was a syringe pump (New Era Pump System Inc., Farmingdale, NY) that was used to drive the syringe plunger to push the hydrogel through a tube, which was connected to the inner nozzle (21 G) of the coaxial assembly. The last component was the collection bath, which was filled with oil or calcium chloride (CaCl_2_) solution as per the precursor hydrogel used. The collection bath was utilized for the collection and crosslinking of microgels. Along with these main components, a magnetic stirrer was used to homogeneously stir the solution in the collection bath. The collection bath was placed on the magnetic stirrer below the coaxial nozzle system, which was fixed with a clamp. Once the setup was built, precursor hydrogel solutions were prepared and microgels were generated and then crosslinked as shown in **Fig. 1B**. The air pressure and flow rate were regulated and optimized to produce different types and sizes of microgels.

### 2.2 Preparation of alginate (Alg) microgels

Alginate microgels were produced by dissolving sodium alginate (Sigma-Aldrich, United Kingdom (UK)) in deionized (DI) water at different concentrations (0.5, 1, and 2% w/v). CaCl_2_ solution was prepared by dissolving 4% (w/v) CaCl_2_ (Sigma-Aldrich, St. Louis, MO) in DI water. The alginate solution was taken into a 10 mL syringe and extruded through the inner part of the coaxial nozzle at flow rates of 10, 50, 100, and 500 μL/min, and air pressure levels of 40, 60, 100, and 180 kPa. After optimization, the flow rate was finalized at 100 μL/min and the air pressure was varied as mentioned before to regulate the size of microgels. The generated microgels were collected in the CaCl_2_ bath for crosslinking. After generating microgels, they were transferred into a 50 mL centrifuge tube and washed thrice using DI water to remove the excess CaCl_2_ and uncrosslinked alginate. The collected microgels were used for further studies.

### 2.3 Preparation of GelMA Microgels

GelMA was synthesized as per previously reported protocol [23], by reacting methacrylic anhydride with gelatin, purified using 12-14 kDa cut off dialysis membrane, and freeze dried. Both fish GelMA and porcine GelMA were synthesized from gelatin from cold fish water skin (Sigma-Aldrich) and type A gelatin from porcine skin (Sigma-Aldrich), respectively. The freeze-dried GelMA was reconstituted to a 10% (w/v) final solution. The 10% GelMA solution from both fish and porcine sources were mixed at a 2:8 ratio, respectively, and then 5% lithium phenyl (2,4,6-trimethylbenzoyl) phosphinate (LAP) (TCI chemicals, OR) was added. The formed GelMA solution was loaded into a 10 mL syringe and extruded at an air pressure of 40, 60, 100, and 180 kPa. Light mineral oil (Sigma-Aldrich, St. Louis, MO) with nonionic surfactant (3% Span 80, Sigma-Aldrich) was used in the collection bath to collect GelMA microgels. A visible light source (GHDO, SOVOL, Shenzhen, China) of 405 nm was used to crosslink GelMA droplets to form microgels. The formed microgels were washed three times with DI water followed by centrifugation to remove the oil and surfactant. The collected microgels were used for further studies.

### 2.4 Preparation of Alg-GelMA microgels

Alg-GelMa microgels were produced using sodium alginate, fish GelMA, and porcine GelMA in deionized (DI) water. A GelMA solution consisting of 10% Fish GelMA:10% Porcine GelMA (2:8) was taken and mixed with 1% alginate solution in a 1:1 ratio. The resulting solution was then used for microgel fabrication. Air pressure levels and flow rates were maintained the same as with the above-mentioned microgels. Since the solution consisted of both GelMA and alginate, photocrosslinking (using 405 nm light) and ionic crosslinking (using CalCl_2_ solution) were used simultaneously to crosslink GelMA and alginate, respectively. These microgels were transferred into 50 mL centrifuge tube and washed thrice using DI water to remove the excess CaCl_2_ and uncrosslinked Alg-GelMA. The collected microgels were used for further studies.

### 2.5 Rate, Shape, Size, and Topography Characterization

To quantify the rate of microgel production, microgels were produced for 2 min and collected in 20 mL of the respective collection bath. The number of microgels were counted using a hemocytometer (ART. No. 1280, Ningbo Hinotek Technology Co., China). To assess the morphology of microgels, circularity was calculated using ImageJ software (National Institute of Health, Bethesda, MD). The value ‘0’ indicated an infinitely elongated polygonal shape and ‘1’ indicated a perfectly circular shape.

Further, laser granulometry with a Mastersizer 3000 (Malvern Panalyticals, Worcester, UK) was carried out to assess the size of microgels, which was quantified as volume weighted mean particle size. The particle size analysis was done for samples produced at a constant flow rate of 100 μL/min and at different air pressure levels as mentioned before to study the dependence of particle size distribution on the air pressure.

The surface topography of prepared microgels was assessed using field emission scanning electron microscopy (FESEM, Apreo S, Thermo Fisher Scientific). The samples were dehydrated using graded ethanol solutions, 20, 50, 60 70, 80, 90, and 100%, sequentially each for 10 min. To ensure the complete removal of water, samples were further dried in a critical point dryer (CPD300, Leica, Wetzlar, Germany). The dehydrated samples were sputter-coated with iridium using a sputter coater (Leica, Wetzlar, Germany) and imaged at an accelerating voltage of 3-5 kV.

### 2.6 Rheological Analysis

Rheological properties of microgels were characterized with a rheometer (MCR 302, Anton Paar, Austria). Triplicates were taken for each sample. The tests were carried out using a 25 mm parallel-plate geometry measuring system maintained at 22 °C with a Peltier temperature control system. The shear thinning behavior of Alg, GelMA, Alg-GelMA microgels was evaluated by performing a flow sweep test, where the shear rate was varied from 0.1 to 100 s^-1^. To evaluate the viscoelasticity of microgels, amplitude sweep tests were performed at a constant frequency of 1 Hz and shear strain was varied from 0.1 to 100%. The frequency sweep tests were executed to study the storage modulus (G’) and loss modulus (G”) within the linear viscoelastic region at a low shear strain of 0.1% to prevent damages to samples and the angular frequency was varied from 0.1 to 100 rad/s. Further, self-healing tests were performed by measuring the viscosity of microgels under alternating low and high shear rates of 0.1 s^-1^ for 60 s and 100 s^-1^ for 10 s, respectively.

### 2.7 Generation of Cell-Encapsulated Microgels

To generate cell-encapsulated microgels, the ACAD platform was prepared under sterile conditions where a 0.22 μm filter was used to sterilize the air and precursor polymer solutions, which were then loaded with green fluorescent protein (GFP^+^) MDA-MD-231 cells and jetted using the ACAD platform. All the work was performed under a Biosafety Level-2 (BSL-2) (LabGard® ES (Energy Saver) Class II, Type A2 Plymouth, MN) safety cabinet. The hydrogel flow rate was maintained constant at 100 μL/min and the air pressure was maintained at 100 kPa for Alg and GelMA, and 60 kPa for Alg-GelMA. The reduced air pressure for Alg-GelMA was a result of their inability to form stable microgels at higher pressure levels. For cell encapsulation in microgels, GFP^+^ MDA-MB-231 cells (5×10^6^ cells/mL) were pre-mixed in the precursor polymer suspensions and formed into microgels. Once cell-laden microgels were generated and crosslinked, they were washed with DI water three times and centrifuged at 2,000 rpm and dispersed into cell media. They were cultured in Dulbecco’s Modified Eagle Medium (DMEM) media supplemented with 10% fetal bovine serum (FBS) (Life Technologies, Grand Island, NY), 1 mM Glutamine (Life Technologies, Carlsbad, CA) and 1% Penicillin-Streptomycin (Life Technologies). MDA-MB-231 cells were used at passages 56 through 64. The formed cell-encapsulating microgels were imaged using a Zeiss Axio Observer microscope (Zeiss, Jena, Germany) after 6 h of culture.

The encapsulation efficiency was assessed using the images, where the total number of cell-encapsulating microgels was dividing by the total number of microgels. Similarly, the number of cells per microgel were counted and the cells per microgel ratio was reported accordingly.

### 2.8 Extrudability and Shape Fidelity Analysis

#### 2.8.1 Hanging Filament Test

Fabricated GelMA microgels were added into 15% porcine GelMA hydrogel in a 1:1 ratio, and then mixed with a vortex mixer. Next, the suspension was centrifuged at 13,000 rpm and the supernatant was removed. This process was repeated twice and the centrifuged GelMA microgels in GelMA hydrogel were used for 3D bioprinting purposes.

Both GelMA microgels alone and GelMA microgels loaded in bulk GelMA were used as a bioink and tested for their maximum filament length to assess their extrusion fidelity for 3D bioprinting. The bioink made of GelMA microgels was filled to a 3 mL syringe barrel and extruded through a needle with the help of controlled pneumatic pressure using an Inkredible^+^ bioprinter (CELLINK, Sweden). The bioink came out in the form of filament and the maximum length of extruded filaments was measured before they broke. The needles used for this test were 25 G (250 μm inner diameter), 22 G (410 μm inner diameter), and 20 G (610 μm inner diameter). The pneumatic pressure was maintained at 10 to 20 kPa for the bioink with GelMA microgels only, and 90 to 120 kPa for the GelMA microgels loaded in GelMA. The videos of filament extrusion were recorded using a Nikon D810 camera with Nikon 105 Micro lens. The frame, at which the filament broke its contact with the needle was taken and the length of the filament was measured using ImageJ.

#### 2.8.2 Filament Fusion Test

To perform the filament fusion test, grids with different pore sizes were 3D bioprinted and the spreading and printability analysis was performed according to the literature. [24]. Briefly, a bi-layered grid with square pores of varying size from 1×1 to 5×5 (mm×mm) was bioprinted on glass slides. Both bioinks, i.e., GelMA microgels and GelMA microgels loaded in GelMA, were extruded through a 22 G needle at a bioprinting speed of 6 mm/s. The former was extruded at a pressure of 15 kPa while the latter was extruded at 100 kPa. The images of bioprinted grids were taken with the Nikon D810 camera with Nikon 105 Micro lens and analyzed using ImageJ. The percentage diffusion (*D*_*f*_) was calculated using Equation 1, which allows one to interpret the degree of bioink spreading after extrusion. The percentage diffusion was ‘0’ in case the bioink did not spread and the actual area became equal to the theoretical area. Here S_*t*_ and S_*a*_ were the theoretical and actual area of the pores, respectively. Further, the printability of the bioinks was calculated using Equation 2, where ‘1’ indicated perfect printability.

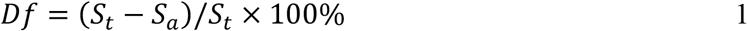

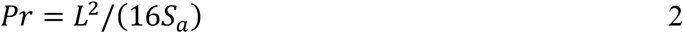

In equation, *L* denoted the perimeter of the actual pore.

### 2.9 3D Bioprinting

GFP^+^ MDA-MB-231 cell loaded GelMA microgels were 3D bioprinted on a build-plate at room temperature using the Inkredible^+^ bioprinter with a 22 G needle. The bioprinting speed and pneumatic pressure were maintained at 6 mm/s and 15 kPa, respectively.

In addition, Alg microgels were explored as a support bath for embedded bioprinting to form complex branched structures. In this regard, Alg microgels were cast into a transparent container. To demonstrate support bath capabilities of Alg microgels, we used embedded bioprinting. For this, xanthan gum was chosen as the bioink material due to its shear thinning properties [25]. A 2% w/v xanthan gum (from *Xanthomonas campestris*, Sigma Aldrich) was prepared in a commercial blender (Magic bullet, Homeland Housewares, CA) for 3 min followed by centrifugation at 4,000 rpm for 5 min. Since the xanthan gum bioink was transparent, oil based red dye with surfactant was added to make it visible during and after bioprinting. Alginate microgels were cast into a transparent container and the red dyed xanthan gum was used to print a 3D structure inside the support bath with the Inkredible^+^ bioprinter. The needle used for embedded bioprinting was 25 G and the bioprinting speed was maintained at 1 mm/s.

### 2.10 Microgels for Scaffolding Application

The formed acellular microgels were seeded with GFP^+^ MDA-MB-231 cells (10 × 10^6^ cells/mL). The distribution of cells in the microgel matrix was assessed after 6 h using a Leica SP8 DIVE multiphoton microscope (Leica Microsystems, Germany) with 16x water immersion lens. Dead cells were assessed by staining with ethidium homodimer-1 (EthD-1, 4 μM) for 30 min in the incubator, which were then imaged using the Zeiss Axio Observer microscope. Images were analyzed using Fiji ImageJ to determine the red fluorescence intensity and the cellular viability was quantified by dividing the fluorescence intensity of dead cells (red) to the combined fluorescent intensity of both dead (red) and live (green) cells.

For evaluation of cell proliferation, MDA-MB-231 cells were seeded with microgels on cell-repellent surface microplates and assessed at Days 1, 3, and 7. Alamar blue dye reduction assay (Invitrogen, USA) was used following the manufacturer’s protocol. Briefly, cell seeded microgels were incubated with 10% (v/v) of the dye for 3 h. After incubation, 100 μL of the culture media was read using a microplate reader (Tecan Infinite 200 Pro, Switzerland) at 570/600 nm (excitation/emission). The results were presented as the normalized value of the dye reduced, which were proportional to the number of viable cells present in microgels.

### 2.11 Statistical Analysis

All data were presented as mean ± standard deviation. Data were analyzed by OriginPro 9.1. Statistical differences were determined with one-way analysis of variance (ANOVA) and Tukey’s post hoc test, and the analysis, if fulfilling the null hypothesis at p ≤ 0.05, was considered as statistically significant (^*^), while at p ≤ 0.01 (^**^) and p ≤ 0.001 (^***^) as highly significant and ‘ns’ represents not significant.

## 3. Results

### 3.1 Fabrication of Microgels and their Physical Characterization

Spherical microgels of Alg, GelMA, and Alg-GelMA were successfully fabricated using the ACAD platform. The precursor polymer solutions were extruded at different pressure levels namely 40, 60, 100, and 180 kPa, and the microgel size and shape was analyzed. The brightfield images of Alg, GelMA, and Alg-GelMA microgels fabricated using above-mentioned pressure levels were shown in **Fig. 2A**. The microgels exhibited spherical morphology and were separated from each other without any visible agglomeration. The efficacy of the ACAD platform was assessed by quantifying the rate of microgel production. The microgel production rate was found to be ∼3,800 microgels/sec for Alg, ∼65,000 microgels/sec for GelMA, and ∼7,000 microgels/sec for Alg-GelMA (**Fig. 2B**). The rate of production for GelMA microgels was significantly higher (p≤0.001) as compared to that for Alg and Alg-GelMA microgels. Further, the circularity of the microgels was determined (**Fig. 2C**). Alg, GelMA and Alg-GelMA microgels showed a circularity distribution ranging from 0.2 to 1, from 0.5 to 1 and from 0.2 to 1 respectively. However, depending on the range, the circularity of GelMA was determined to be significantly higher than that for Alg (p≤0.01) and Alg-GelMA (p≤0.001). Overall, the majority of Alg and GelMA microgels exhibited a circularity score of ∼1, indicating a perfect circular shape. Alg-GelMA microgels were large and less circular, which was also confirmed by SEM micrographs (**Fig. S1**).

**Fig. 2:**
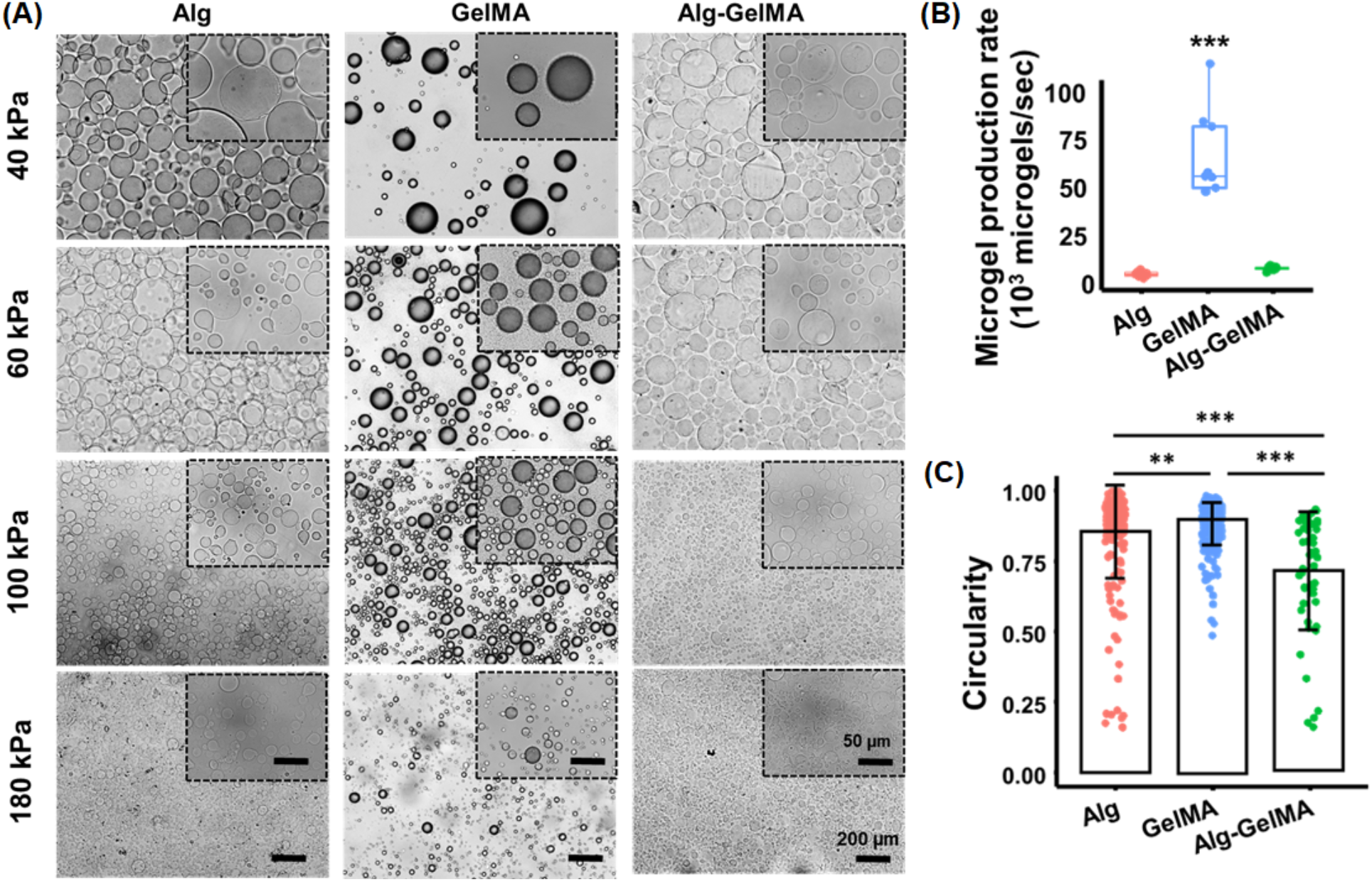
Morphological assessment and physical characterization of acellular microgels. (A) Microscopic images of Alg, GelMA, and Alg-GelMA microgels at different air pressure levels for visualization and comparison of shape and size (scale bar: 200 μm); images in the inset show microgels at a higher magnification (scale bar: 50 μm). (B) Microgel production rate (*n*=9). (C) Quantitative assessment of circularity for the generated microgels (different number of microgels were considered for Alg (*n*=222), GelMA (*n*=252) and Alg-GelMA (*n*=54) (p^**^<0.01, p^***^<0.001).

The particle size distribution was assessed using a particle size analyzer and the plots were presented in **Figs. 3A-C**. Microgels with sizes ranging from 10 to 1000 μm were obtained based on varying air pressure levels. The microgel size decreased with increasing pressure levels. It was noted that the particle size maxima of the microgels decreased from 186 to 46 μm for Alg, 350 to 86 μm for GelMA, and 454 to 76 μm for Alg-GelMA with an increase in pressure from 40 to 180 kPa (**Figs. 3D-F**). Alg microgels were relatively smaller in size followed by GelMA and Alg-GelMA counterparts at each pressure levels. After generating all microgels at different pressure levels, we noticed the generation of larger microgels at lower pressure levels (40, 60 kPa), which were not suitable for bioprinting applications while smaller size microgels were observed at a high-pressure level (180 kPa) that might impede cell encapsulation. Therefore, we preferred 100 kPa air pressure for Alg and GelMA microgels. For Alg-GelMA microgels, we used 60 kPa since we did not get stable microgels at 100 kPa.

**Fig. 3:**
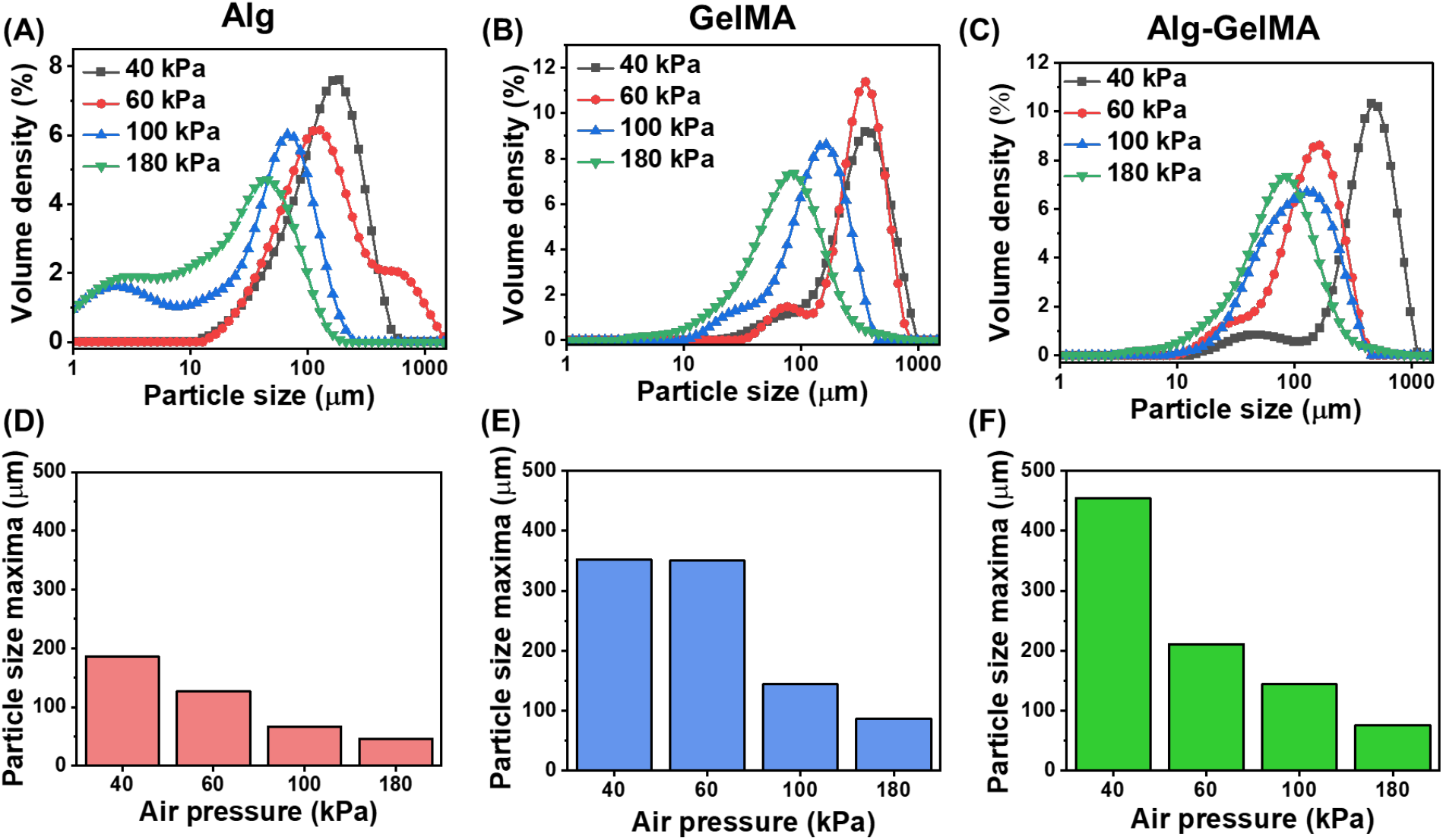
Particle size distribution analysis of microgels under different air pressure levels and a fixed flow rate. The volume density distribution profiles and particle size maxima of (A, D) Alg, (B, E) GelMA, (C, F) Alg-GelMA microgels.

Rheological assessment of microgels was performed to confirm that they were appropriate for specific applications such as an extrudable bioink or a support bath for 3D bioprinting. The shear thinning behavior of microgels was studied with a flow sweep test by measuring their viscosity at varying shear rate ranging from 0.1 to 100 s^-1^. All groups possessed shear thinning behavior and the viscosity decreased with increasing shear rate. The viscosity of Alg-GelMA was the highest up to a shear rate of 10 s^-1^ and decreased rapidly after that and became the lowest after 20 s^-1^ (**Fig. 4A**). The viscosities of Alg and GelMA microgels were nearly similar in the entire test range.

**Fig. 4:**
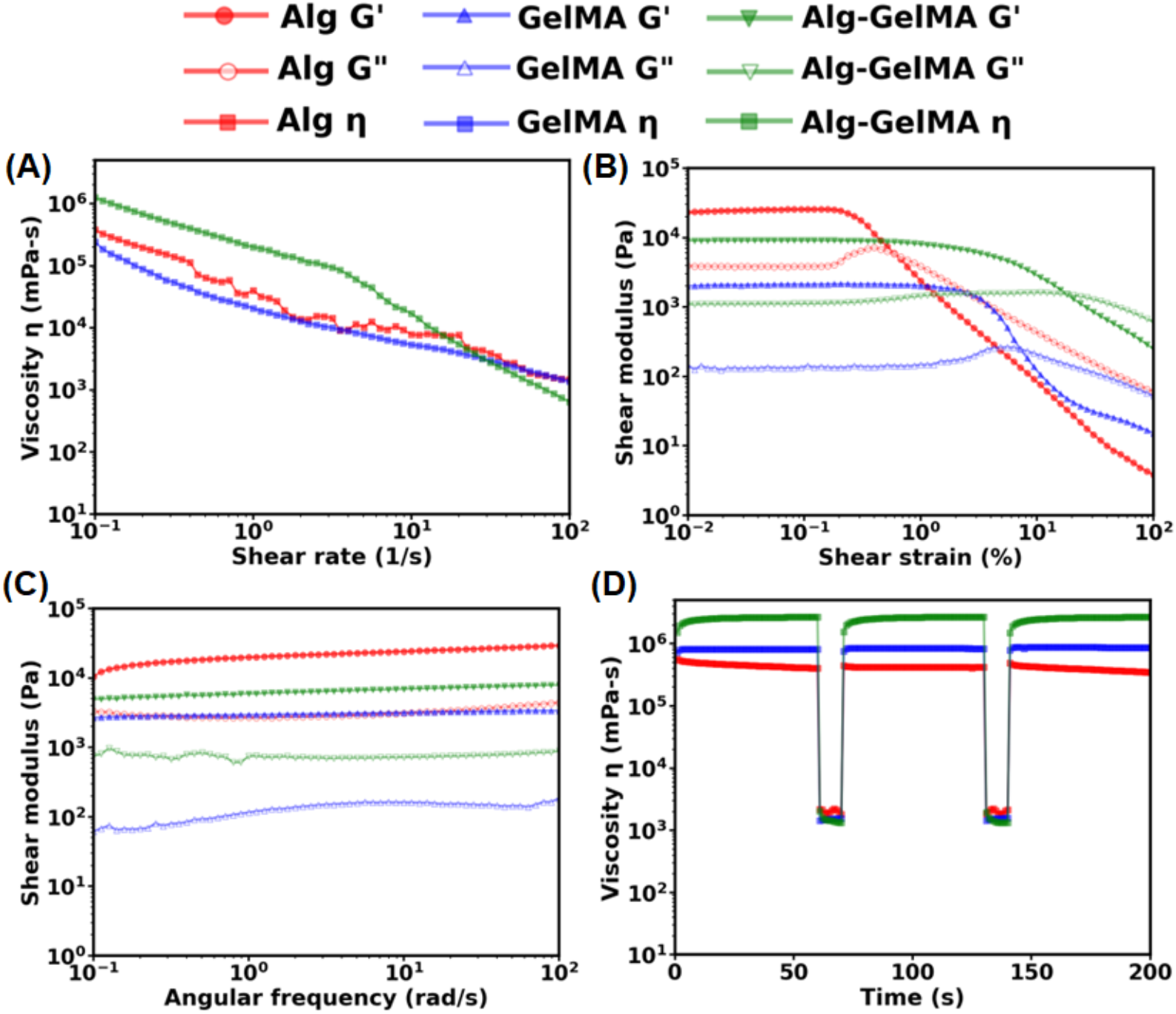
Rheological characterization of microgels. (A) Flow curves showing viscosity as a function of shear rate ranging from 0.1 to 100 s^−1^. (B) Amplitude sweep test to measure shear modulus of microgels at a strain ranging from 0.01 to 100%. (C) Frequency sweep test of microgels in the linear viscoelastic region at an angular frequency ranging from 0.1 to 100 rad s^−1^. (D) Self-healing tests to assess the recovery of microgels after applying a high shear stress. The viscosity was measured at five intervals at alternating shear rates of 0.1 s^−1^ for 60 s and 100 s^−1^ for 10 s.

The amplitude sweep test demonstrated the transition of microgels from an elastic state (G’ > G”) to viscous state (G” > G’) as strain was increased from 0.01 to 100 %. All the samples initially showed higher G’ than G” stating that microgels behaved as elastic materials under low values of shear strain and transitioned to a viscous state as the shear strain increased. It also illustrated that the storage modulus of Alg, GelMA, Alg-GelMA microgels was ∼250, ∼2, and ∼10 kPa, respectively, at low shear strain values until they reached their yield stress point, which is defined as the point where the microgels transitioned from elastic to viscous state (**Fig. 4B**). The percentage strain at which the material yields was found to be the highest for Alg-GelMA (20%), followed by GelMA (7%) and Alg (0.5%). The frequency sweep test demonstrated G’ as superior to G” for all microgels indicating them as elastic materials at low shear strain 0.01% and over the entire range of angular frequency 0.1-100 rad/s (**Fig. 4C**).

The self-healing test was also performed to assess the recovery of microgels after deformation that occurs because of a high shear rate particularly when using them as a support bath. The self-healing properties were studied by varying the shear rate from low (0.1 s^-1^) to high (100 s^-1^) alternatively in cycles. The low shear rate was applied to mimic the static condition while the high shear rate was applied to mimic the stress generated during movement of a needle in a support bath. The healing behavior was observed by studying the difference in the viscosities before and after applying the high shear rate. As shown in **Fig. 4D**, it was found that all microgel types recovered completely, indicating their self-healing behavior. The change in the viscosity was more substantial for Alg-GelMA microgels compared to their Alg and GelMA counterparts.

### 3.2 Applications of Alg, GelMA and Alg-GelMA microgels

To exemplify the utility of fabricated microgels, we demonstrated three distinct applications including cell encapsulation, bioprinting (both direct extrusion of microgels and their use as a support bath) and scaffolding.

In the first application, we demonstrated cell encapsulation. Upon combining GFP^+^ MDA-MB-231 cells with the precursor polymer solution, cell-encapsulated microgels were produced as shown in **Figs. 5A, S2**. The encapsulation efficiency of MDA-MB-231 cells in microgels was found to be ∼75% for all three microgel types (**Fig. 5B**). Further, the number of cells per microgel ratio was ∼3.2 for Alg, ∼1.5 for GelMA, and ∼2.7 for Alg-GelMA microgels (**Fig. 5C**).

**Fig. 5:**
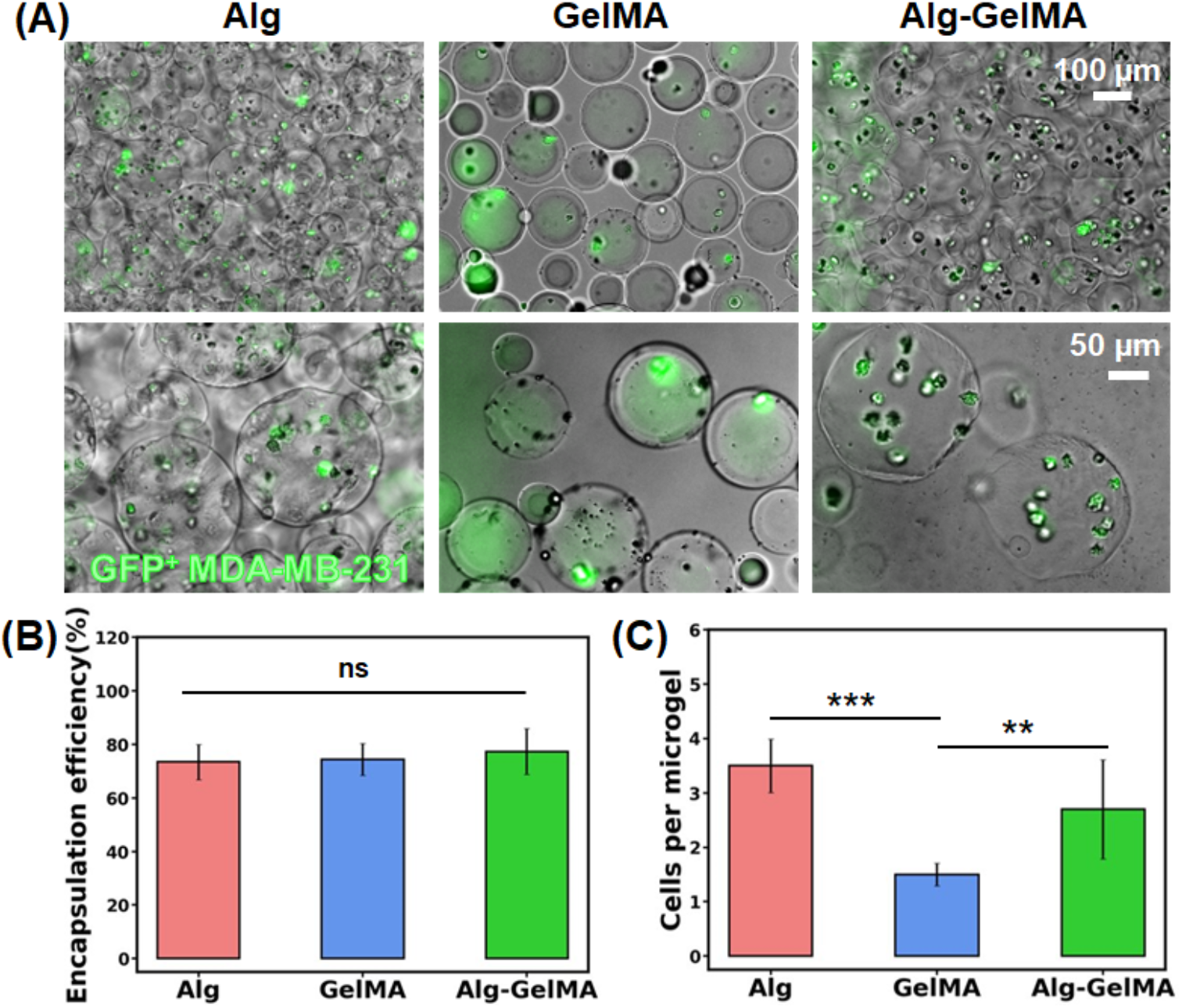
Cell encapsulation. (A) Fluorescence microscopy images showing GFP^+^ MDA-MB-231 cells encapsulated in microgels. (B) Cell encapsulation efficiency and (C) the number of cells per microgel in Alg, GelMA, and Alg-GelMA microgels (*n*=5, p^**^<0.01, p^***^<0.001, ns – not significant).

In the second application, microgels were used as a bioink and a support bath for extrusion-based bioprinting. Based on rheological characteristics and preliminary extrusion testing, only GelMA microgels were taken further for their assessment as a printable bioink because Alg and Alg-GelMA microgels alone were not extrudable as they showed water phase separation, did not form filaments, and caused nozzle clogging. The characterization tests for the extrusion of GelMA microgels revealed acceptable printability and shape fidelity, which were further improved after loading GelMA microgels in bulk GelMA. The hanging filament test showed that GelMA microgels had a capacity to form filaments, which is necessary for bioprinting as shown in **Figs. 6A, C**. Further, the addition of bulk GelMA into GelMA microgels significantly increased (p≤0.001) the maximum length of hanging filaments thus increasing the robustness of the bioink for extrusion (**Figs. 6B, C**). It was also found that 22 G needles formed thinner filaments than 20 G needles while 25 G needles tended to clog. Therefore, 22 G needles were used for all characterization experiments pertaining to extrusion-based bioprinting.

**Fig. 6:**
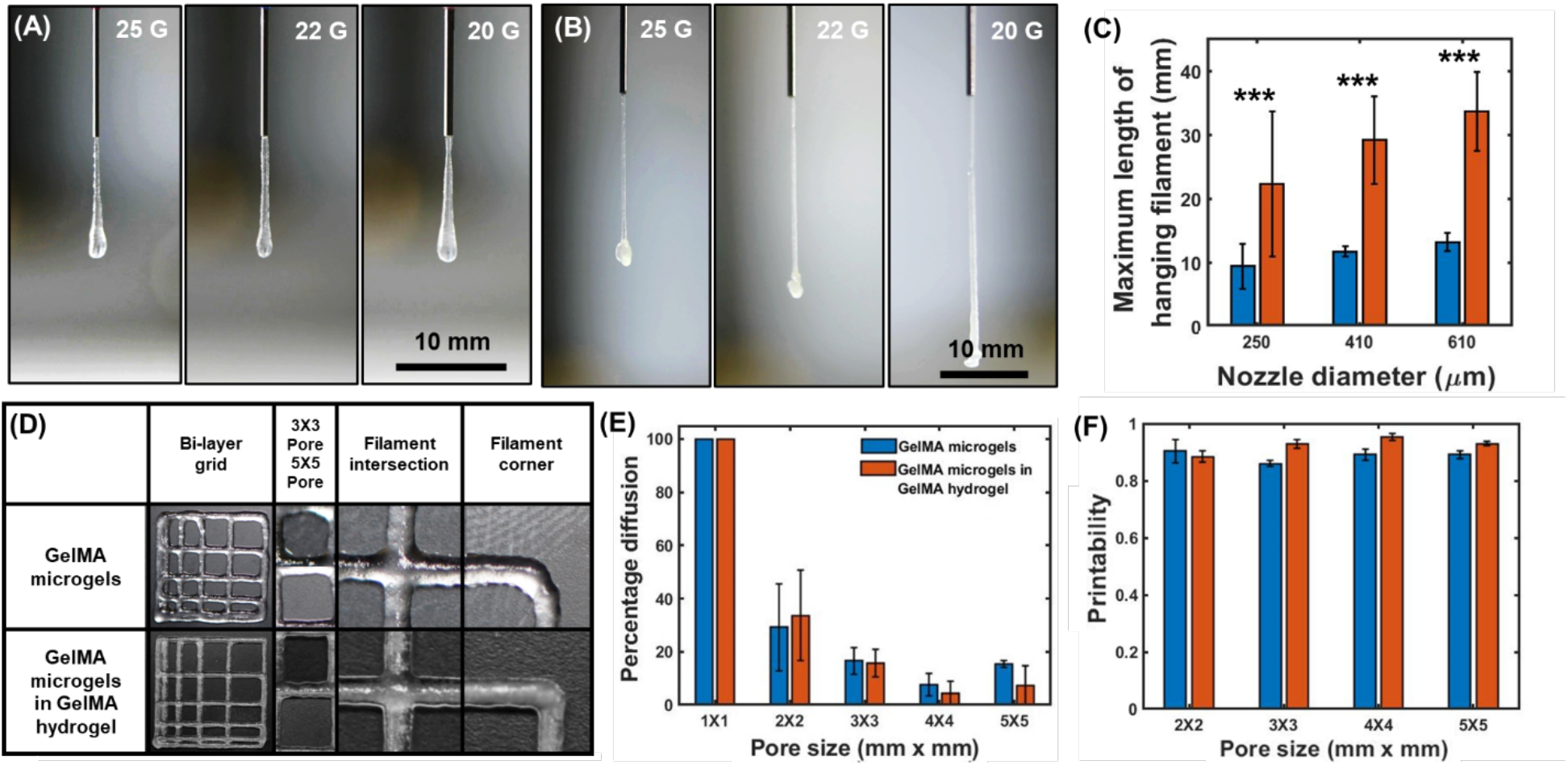
Filament length, fusion, and printability tests for multiple bioinks including GelMA microgels and GelMA microgels loaded in bulk GelMA. Representative images of hanging filaments for (A) GelMA microgels, (B) GelMA microgels loaded in a bulk GelMA hydrogel, (C) quantification of the maximum length of hanging filaments for the two bioinks. (D) 3D Bioprinted grid structures of various known dimensions. (E) The percentage diffusion of bioprinted filaments, and (F) printability of both bioinks (*n*=3, p^***^<0.001).

The filament fusion test was then performed by 3D bioprinting grid structures of different dimensions. The results revealed that the percentage diffusion was greater than 0, which was expected as the filaments slightly deformed and diffused after deposition. Pores of 1 mm × 1 mm grid structures were completely closed resulting in a pore area of 0. Therefore, the diffusion percentage was taken as 100%. However, the percentage diffusion was less than 40% for the 2 mm × 2 mm grid structures, which decreased further under 20% for larger pores as shown in **Figs. 6D, E**. The printability of both bioinks was determined to be greater than 0.8 (**Fig. 6F**) indicating that they have potential to be utilized as an extrudable bioink for 3D bioprinting applications.

After validating that GelMA microgels were printable on their own (**Figs. 7A, B**) or when loaded in bulk GelMA (**Figs. 7D, E**), GFP^+^ MDA-MB-231 cells were successfully encapsulated in GelMA microgels and bioprinted as shown in **Figs. 7C, S2**. Similar bilayer grid structures were successfully fabricated in centimeter scale and cell encapsulating GelMA microgels was bioprinted in the shape of ‘PSU’ (**Figs. 7G-H**), which confirms the printability and scalability of these constructs made of cell encapsulating microgels. Further, embedded bioprinting within Alg microgels was also successfully executed, where Xanthan gum was bioprinted into a branched network to demonstrate the potential of microgels as a support bath (**Fig. 7I**).

**Fig. 7:**
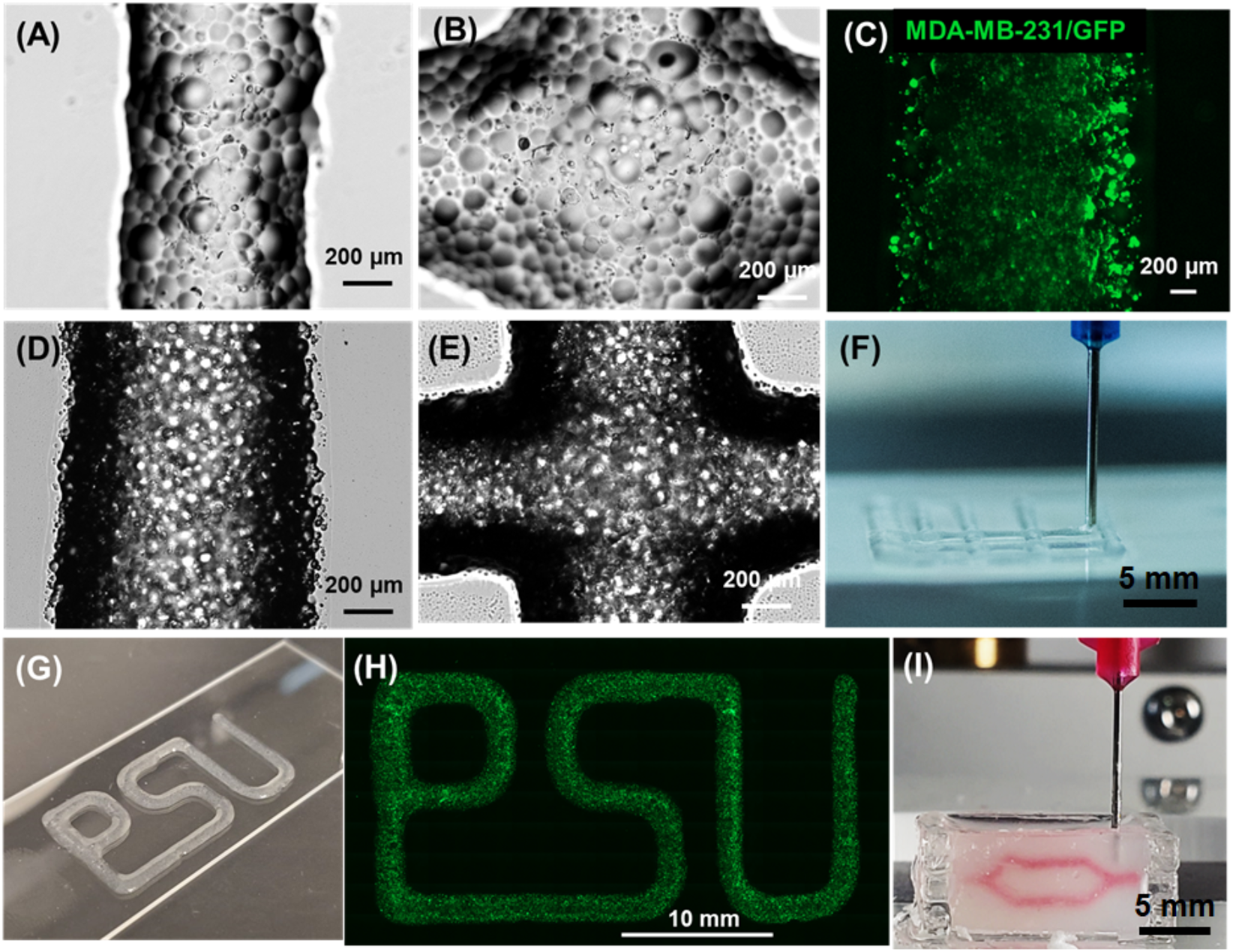
Visual demonstration of bioprinted constructs. Brightfield images showing (A) bioprinted filaments and (B) intersections made of GelMA microgels; (scale: 200 μm). (C) Micrograph of GFP^+^ MDA-MB-231 cells encapsulated in bioprinted GelMA microgels (scale: 200 μm). Brightfield images of (D) bioprinted filaments, and (E) intersections of GelMA microgels loaded in GelMA. (F) 3D Bioprinting of bilayer grid structures. (G) A photograph and (H) a fluorescent image illustrating cell encapsulating GelMA microgels bioprinted in the shape of ‘PSU,’ the abbreviation of Penn State University. (I) Embedded bioprinting of a branched structure inside a support bath made of Alg microgels.

In the last application, we used Alg, GelMA and Alg-GelMA microgels as scaffolds and seeded GFP^+^ MDA-MB-231 cells on them. We evaluate the biocompatibility of microgels by assessing the viability and proliferation of MDA-MB-231 cells. Cells seeded on microgels showed homogenous distribution after 6 h of seeding, as shown in **Fig. 8**. They could evenly surround the microgels and fill the voids between microgels in 3D (**Video S1**). For the viability assessment, GFP^+^ cells were stained with EthD-1 to detect the dead cells as indicated by the red fluorescence signal (**Figs. 8C-D, S3**). MDA-MB-231 cells exhibited a viability of ∼98% on Day 1, ∼95% on Day 3, and ∼90% on Day 7 (**Fig. 8E**), which readily adhered to and proliferated among the microgels in interstitial void spaces. In addition, the AlamarBlue assay showed ∼1.15 and ∼1.2-fold increase in the metabolic activity for cells on Days 3 and 7, respectively, for Alg microgels, ∼1.2 and ∼1.4-fold increase in the metabolic activity of cells on Days 3 and 7, respectively, for GelMA microgels; and ∼1.1 and ∼1.3-fold increase in the metabolic activity for cells on Days 3 and 7, respectively, for Alg-GelMA microgels (**Fig. 8F**). On Day 7, MDA-MB-231 cells seeded on GelMA microgels showed significantly higher proliferation as compared to cells seeded on Alg microgels (p≤0.01).

**Fig. 8:**
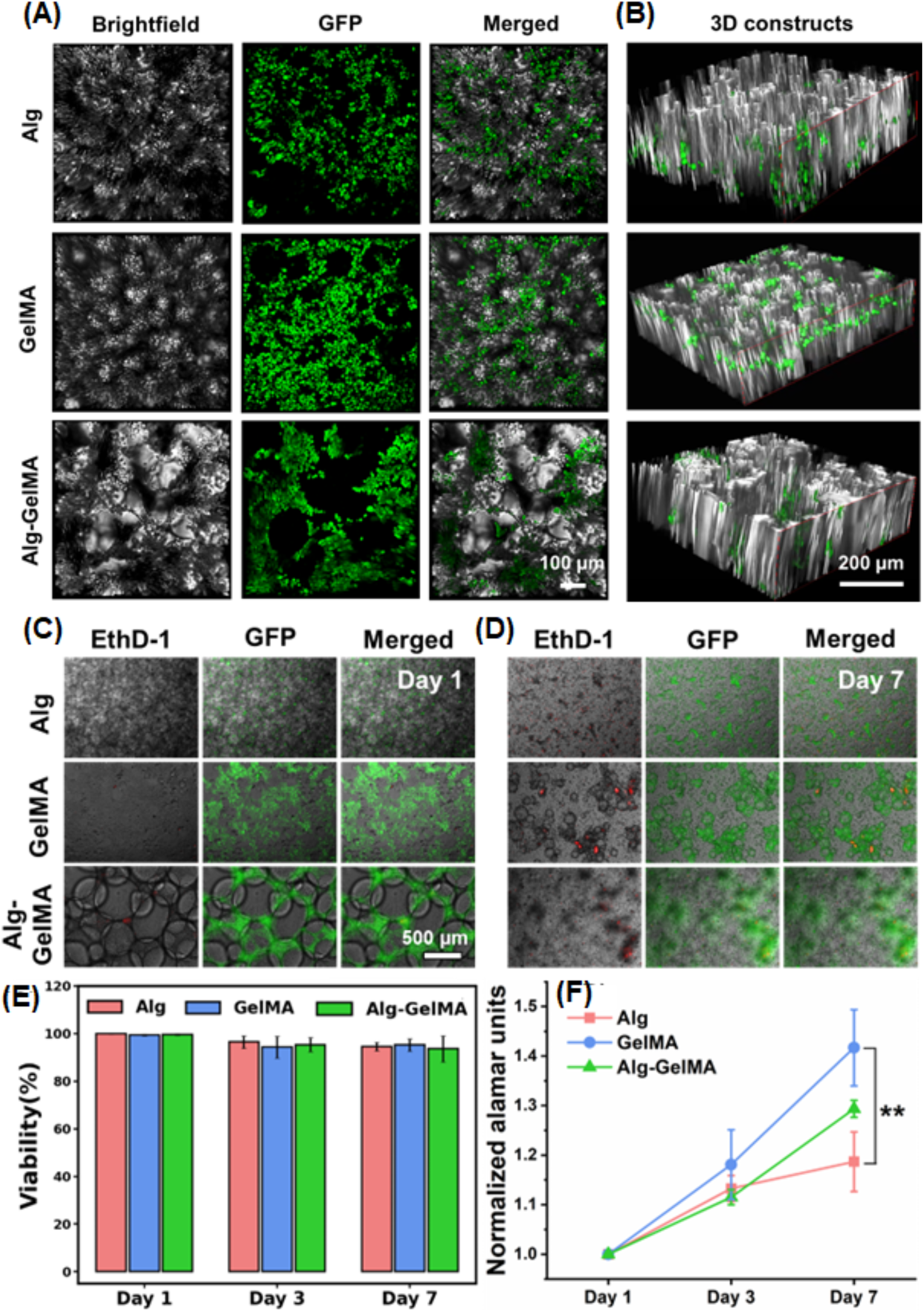
Microscopic images of GFP^+^ MDA-MB-231 cells seeded on microgels. (A) Images showing microgels under brightfield, fluorescence, and merged, and (B) the distribution of cells in 3D. (C) Representative images of GFP^+^ seeded cells (green) and dead cells (red) on Days 1 and 7. (E) Quantification of cellular viability for microgels at Days 1, 3, and 7. (D) Proliferation of cells over a week culture (*n*=3, p^**^<0.01).

## 4. Discussions

Microgels are swollen macromolecular networks that differ significantly from typical colloids, such as micelles, vesicles, flexible macromolecules, and rigid nanoparticles. This is mainly because of the presence of a crosslinker that holds the microgel together and provides structural integrity and control over their characteristics [26]. The microgel fabrication technique described here is advantageous for cell encapsulation, bioprinting and scaffolding since it opens the possibility to develop biomaterials on a scale of tens to hundreds of microns in a high-throughput manner. It is challenging to work with bulk self-assembled polymeric gels in in-vitro culture or after in-vivo implantation without generating cracks and heterogeneities. In consequence, it would be anticipated that these heterogeneities will predominate the movement of media or physiological fluids throughout and within the materials. Here comes the role of the assembled microgels as they can modify their volume and shape in response to external stimuli (such as pressure, light, electrochemical stimulus, temperature, and pH), which enable them to reversibly tune their physicochemical characteristics.

In the current study, we successfully produced microgels of varied sizes by changing the air pressure flowing through the outer channel of the co-axial nozzle system while the polymer solution flows through the core channel. This co-axial nozzle system resulted in the formation of droplets via the mechanism of atomization. These droplets immediately get crosslinked when they reached the collector and thus formed stable microgels. The flow rate of the polymer solution was optimized to be 100 μL/min. In our preliminary studies, we observed the formation of large-sized microgels when the flow rate was in the range of 10-50 μL/min. These large microgels tended to break easily during bioprinting. On the other hand, microgels were extremely small when we used higher flow rates (200 – 500 μL/min). These microgels could not be used for cellular encapsulation. Thus, to obtain stable microgels in a suitable size range for bioink development and cell encapsulation, a constant flow rate of 100 μL was used. Herein, Alg microgels were formed by flowing alginate through the core nozzle and microgels were collected in the CaCl_2_ bath. In an early study, Pravinata et al. produced calcium alginate microgel particles (∼100 nm) using a jet homogenizer for highly turbulent mixing of two liquid streams of sodium alginate and CaCl_2_ solution [27].

For GelMA microgels, the GelMA solution containing the LAP photoinitiator flowed through the inner nozzle and droplets were collected in an oil bath, where microgels were formed due to water-in-oil emulsion and crosslinked using the 405 nm light exposure. In our study, it is pertinent to note that GelMA from fish and porcine sources were used in combination. This was because we needed continuous flow of GelMA, and the porcine GelMA exhibited thermal crosslinking at room temperature that made it hard to flow while fish GelMA was liquid at room temperature and weak in terms of crosslinking. Therefore, we formulated a composite hydrogel by mixing them to get the required flow properties at room temperature. Nichol et al. fabricated cell-laden GelMA microgels down to 100 μm resolution with the fidelity and robustness by assembling microgels at the surfactant-containing oil–water interface followed by secondary crosslinking via ultraviolet (UV) light using Irgacure 2959 photoinitiator [28]. Further, Alg-GelMA microgels were produced in this study by flowing a LAP-containing blend of alginate and GelMA solution through the nozzle and collecting it in the CaCl_2_ bath and exposing to 405 nm light to accomplish dual crosslinking. To the best of our knowledge, Alg-GelMA microgels have not been reported previously in the literature.

Herein, we studied the effect of change in air pressure on the size of formed microgels. For this, different air pressure levels ranging from 40 to 180 kPa were investigated and the results indicated that at >100 kPa, the microgel size was smaller than desired, on the other hand, they were too large at ≤60 kPa. Therefore, we preferred 100 kPa for further studies to get microgels at a consistent size, which was required for cellular encapsulation and bioprinting. In a study, Akbari et al. developed a microfluidic multiplier droplet maker capable of producing ∼50 μm microgels with a single channel throughput of up to 75 grams per day for 8 μm poly(ethylene glycol diacrylate) microgels at a drop generation frequency of 3.1 MHz. When the oil phase was driven through the micropillar array, shear forces from the oil phase caused the continuous stream of cell suspension to break up into droplets [8]. Here, we used air as our continuous phase and found that increasing the air pressure resulted in decreased microgel size. Recently, Li et al. reported a high-throughput approach, where they used a multi-channel rotating system to generate alginate microgels ranging from 100-1,000 μm in size [29], but the reported setup was very complex as compared to the one reported in our study..

The formed microgels were then characterized for the rate of microgel production and circularity. The results showed that ∼3,800, ∼65,000, and ∼7,000 microgels per sec were produced for Alg, GelMA, and Alg-GelMA, respectively. For comparison with the literature, Morimoto et al. reported mass production of calcium-alginate particles with a rate around 171 particles per sec with the size range of 100-300 μm [30]. Additionally, the circularity assessment in our study indicated that the formed microgels had a wide range of circularity. GelMA microgels showed the highest circularity, closely followed by Alg microgels. Alg-GelMA microgels showed the least circularity, which might be due to the improper blending of the two polymers and their incomplete crosslinking especially in the core of these large microgels. Therefore, the main factors determining the final size of microgels include the polymer flow rate, air pressure at the co-axial nozzle, and microgel composition.

Further, a systematic rheological analysis was carried out to examine the three types of microgels and evaluate their applicability as an extrudable bioink for bioprinting or as a support bath for embedded bioprinting. The decrease in the viscosity of GelMA microgels upon increasing the shear stress signifies that the applied stress overcame the inter-microgel interactions of GelMA. It led microgels to move relative to each other, and after crossing the yield point, microgels were no longer elastic and behaved like a viscous fluid. Further, the amplitude sweep test also validated that GelMA microgels behaved as a yield stress gel, and their mechanical regime transformed from elastic to viscous nature near 7% strain. This shear thinning effect is crucial for uniform extrusion through a nozzle during bioprinting [31]. Furthermore, an equally important requirement is the recovery of microgels to their original arrangement on the build plate after exposure to a higher stress inside the nozzle. This is called self-healing capability and is needed for bioprinted construct to remain intact. GelMA microgels demonstrated self-healing capability highlighting the fact that they rearranged and packed themselves again immediately after the removal of higher shear rates. Alg microgels also showed shear thinning behavior as their viscosity decreased with increasing shear stress. Since we utilized Alg microgels as a support bath for bioprinting, this shear thinning functionality was crucial as it allowed smooth movement of the needle inside the bath without breaking it. Further, the self-healing capability of these microgels resulted in the quick recovery into their original shape after the removal of higher shear stress. Therefore, Alg microgels were successfully used as an appropriate support bath for embedded bioprinting. Like GelMA and Alg microgels, Alg-GelMA microgels also exhibited shear thinning and self-healing properties. Therefore, they have potential to be used as an extrudable bioink as well as a support bath.

Although rheological data showed promising results for microgels as a yield stress bioink and a support bath, bioprinting experiments should be performed to verify its applicability in bioprinting. During bioprinting, there can be additional factors like gravity, surface tension, clogging, and drying of bioink, which play a crucial role in extrusion and can alter the final extrusion outcomes [32]. During our study, it was noticed that GelMA microgels were suitable for bioprinting only up to a few layers and therefore not recommended for developing hanging constructs. However, when GelMA microgels were loaded in bulk GelMA, the composite bioink gained the ability to facilitate 3D self-standing constructs as validated with the significantly longer filaments shown in **Fig. 6**. Alg microgels showed their potential as a support bath owing to their promising yield stress properties [26, 27]; however, further optimization may be needed to improve the support bath properties for embedded bioprinting depending on the bioink utilized.

For the scaffolding application, GFP^+^ MDA-MB-231 cells were seeded on microgels, which showed immediate penetration after their topical seeding on microgels based on the acquired 3D z-stacked images. Cell penetration into microgels was dictated by the pore size distribution, interconnectivity, and their packing. Herein, no separate jamming of microgels was performed except for the centrifugation during the processing and washing of microgels. This contributes towards a facile and rapid approach of microgel production. The cellular viability for all microgels was found to be ∼90-98 over a period of seven days indicating the biocompatibility of all three microgel types. However, cell proliferation with GelMA microgels was shown to be significantly higher as compared Alg. This is plausibly due to the inherent material properties of polymers. Alg is negatively charged, and as cell surface is also negatively charged, it limits the ability of cells to bind, and this has been explored for alginate’s application in immune-isolation of pancreatic islets [28, 29], among other applications. GelMA, on the other hand, is a biocompatible material with tunable physical characteristics. It has native extracellular matrix (ECM) like integrin-binding motifs and matrix metalloproteinase sensitive groups, which make it suitable for cell attachment and proliferation [36]. Moreover, its polymerization using LAP allows crosslinking in visible or near UV range (405 nm) making it safer for cells. The limitations of Alg in terms of bioactivity could be overcome by combining it with GelMA. The concentration and crosslinking methods of the two different hydrogel-precursor solutions can be tuned to enable a higher resolution of cell-laden microgels to recreate the ideal microenvironment, cell spreading, and organization. The formed Alg-GelMA microgels supported higher cell proliferation as compared to Alg alone.

The microgel fabrication technique described here is advantageous for cell encapsulation, 3D bioprinting, and scaffolding applications since it helps define microgels on a scale of tens to hundreds of microns. Encapsulating cells in microgels allows their easy transfer from one container to another without inducing major stress on them. These microgels could potentially encapsulate different cell types or using various microgel types encapsulating different cell types forming multilayered structures. The application of the produced microgels in the current study could be extended to encapsulate single cells. Current cell encapsulation methods typically produce high polymer-to-cell ratios and lack control over the hydrogel’s mechanical properties [30]. Mao et al. reported a microfluidic-based method for encapsulating single cells in a ∼6 μm layer of alginate that increases the proportion of cell-containing microgels by a factor of ten, with encapsulation efficiencies over 90% [31].

To further advance the presented technology, detailed investigations are required for optimization of certain aspects including the ability to form homogenous microgels of same size. Depending on the application, the size could be tuned or methods such as filtering out microgels of selective size could be opted. Controlling the size could also be extended towards achieving single cell encapsulated microgels. The size and shape of Alg-GelMA microgels could not be tuned enough as stable microgels were not formed at pressure above 80-100 kPa. Thus, its application is limited to strategies involving larger microgels. Towards improved usability of microgels, their cell encapsulation efficiency and the number of cells per microgel ratio could be further controlled by tuning the cell concentration used in the precursor solution and the size of microgels. However, the size of microgels is linked to the air pressure and this has an upper cap as cells might disintegrate at higher air pressure levels.

## 5. Conclusion

In this study, Alg, GelMA and Alg-GelMA microgels were produced using an air-assisted co-axial device in a continuous and high-throughput manner and their utilities were exemplified in multiple applications. Herein, we showed a much higher rate of microgel production particularly for GelMA microgels, where microgel size depended on the air pressure. The results indicate that the proposed Alg microgels have the potential to be used as a prospective support bath material while GelMA microgels have potential for direct extrusion both on their own or loaded with bulk GelMA. Microgels supported cellular attachment, viability, and proliferation, particularly GelMA microgels, and showed high encapsulation efficiency with the capability to encapsulate single to multiple cells. Overall, this study illustrates a facile, affordable, and rapid method for microgel production which can be tuned further for various potential applications including encapsulation and delivery of cells, drugs, or other therapeutic and bioactive molecules and to fabricate clinically-relevant cell-laden structures via 3D bioprinting.

## Acknowledgments

This work has been supported by National Science Foundation Award 1914885, National Institute of Dental and Craniofacial Research Award R01DE028614, National Institute of Allergy and Infectious Diseases Award U19AI142733, 2236 CoCirculation2 of TUBITAK award 121C359. We thank Dr. Danny Welch, from University of Kansas, Kansas City, USA for gifting GFP^+^ MDA-MB-231 metastatic breast cancer cells used in the study. The opinions, interpretations, conclusions, and recommendations are those of the author and are not necessarily endorsed by National Science Foundation, National Institute of Dental and Craniofacial Research, National Institute of Allergy and Infectious Diseases Award and TUBITAK.

## Data availability statement

All data that support the findings of this study are included within the article (and any supplementary files). Additional data related to this paper may be requested from the authors.

## Competing interests

I.T.O. has an equity stake in Biolife4D and is a member of the scientific advisory board for Biolife4D and Brinter. Other authors confirm that there are no known conflicts of interest associated with this publication and there has been no significant financial support for this work that could have influenced its outcome.

