## Supplementary Information for "High-throughput microgel biofabrication via air-assisted co-axial jetting for cell encapsulation, 3D bioprinting, and scaffolding applications"

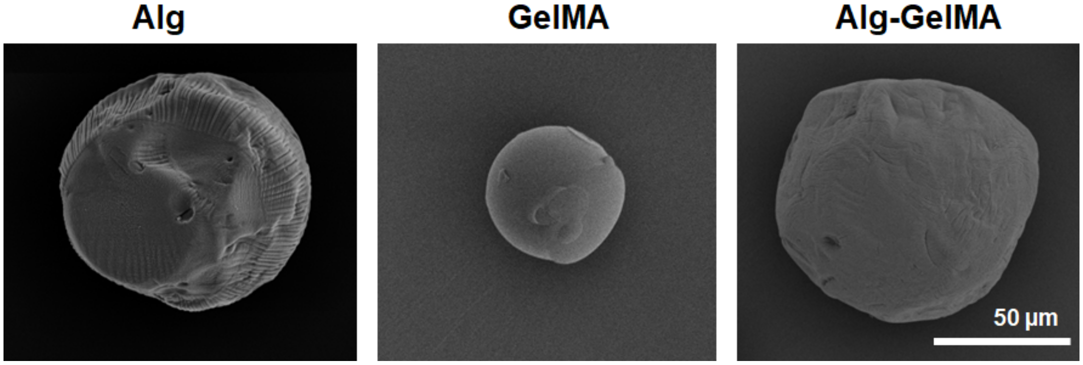


**Fig. S1:** SEM images of Alg, GelMA and Alg-GelMA microgel showing their morphology.

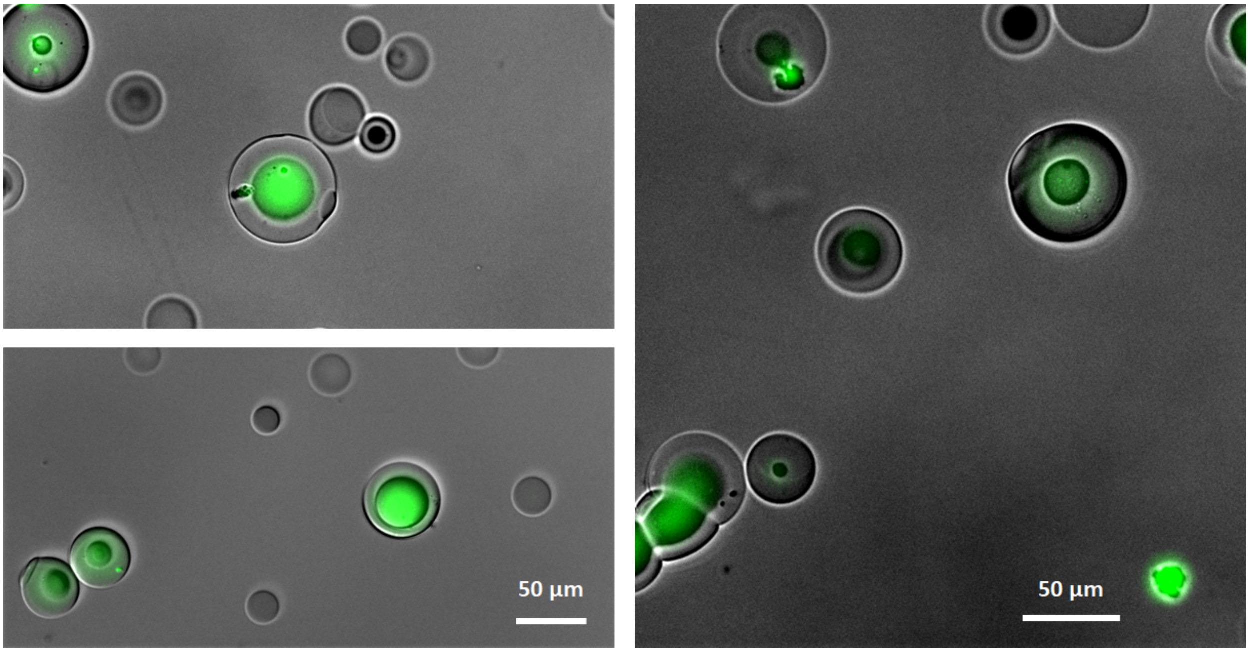


**Fig. S2:** Fluorescence microscopy images showing $GFP^{+}$ MDA-MB-231 cells encapsulated in GelMA microgels.

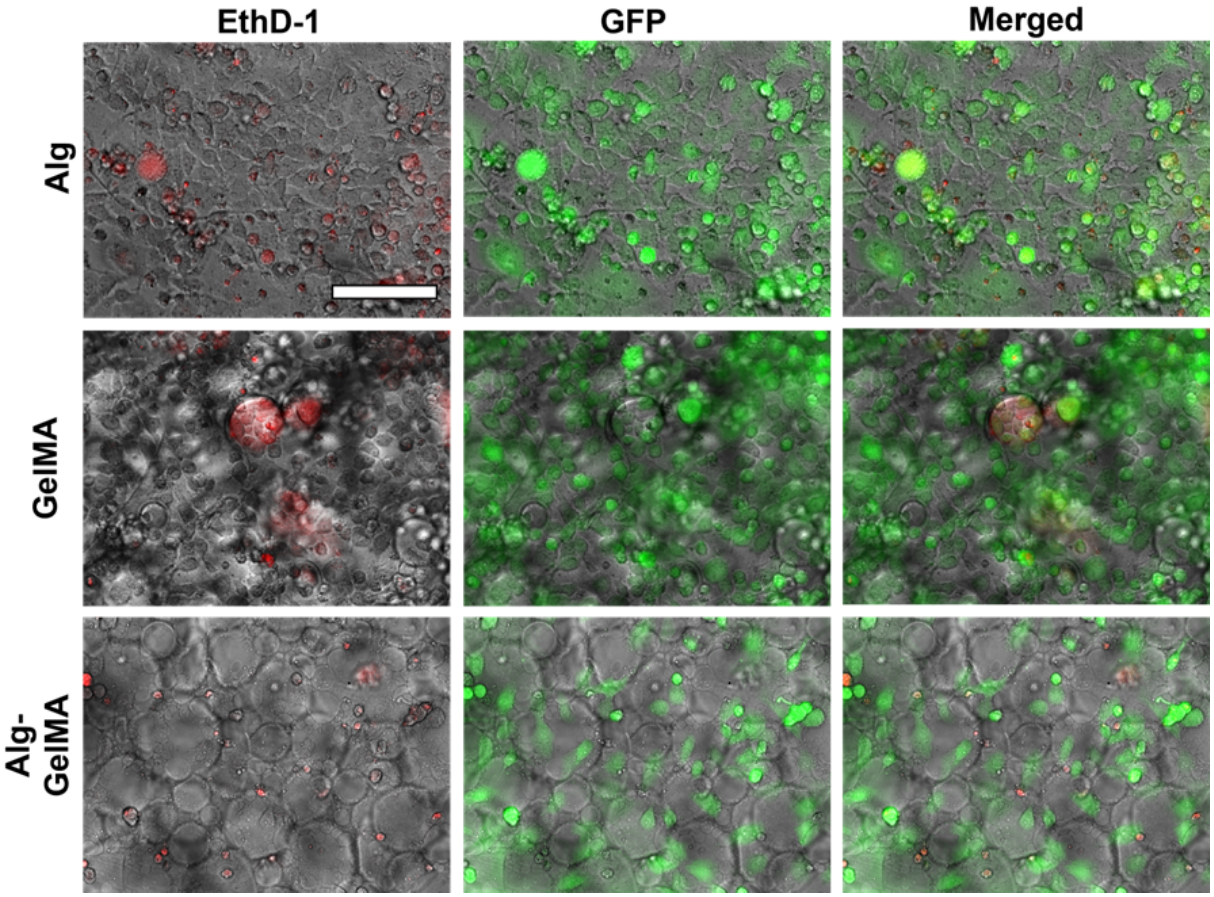


**Fig. S3:** Representative images of GFP$^+$ seeded cells (green) and dead cells (red) on Day 3 (scale bar: 500 μm).

**Video captions:**

**Supplementary Video S1:** 3D Reconstructed multiphoton microscope images of Alg, GelMA, and Alg-GelMA microgels showing the distribution of cells (green) in the microgel matrix.
